# Templated seeding induces the formation of Alzheimer paired helical tau filaments in human brain organoids

**DOI:** 10.64898/2026.09.30.755295

**Authors:** Mehtap Bacioglu, Taxiarchis Katsinelos, Manuel Schweighauser, Filomena Gallo, Karin H. Müller, Rafailia Christou, Janine Brandes, Sofia Lövestam, Sally A. Cowley, William A. McEwan, Sjors H.W. Scheres, Michel Goedert, Maria Grazia Spillantini

## Abstract

The prion concept postulates that seeded aggregation results in the recruitment of soluble protein monomers and their assembly into amyloid filaments with the same structures as those of the seeds. Here we show that this concept applies to tau filaments formed in cortical organoids obtained from human iPSCs of control subjects. Tau seeds were extracted from the frontal cortex of individuals with Alzheimer’s disease. When added to human cortical organoids expressing endogenous levels of three-repeat tau, they gave rise to filaments of three-repeat tau whose formation required endogenous tau expression. By cryo-EM, these seeded tau filaments were identical to the paired helical filaments of the seeds. This novel finding demonstrates the faithful templated seeding of tau in human nerve cells expressing endogenous levels of tau and opens the way to studying the mechanisms of paired helical filament formation and their downstream effects in a human system amenable to genetic modification.

## INTRODUCTION

Alzheimer’s disease (AD) is defined by the presence of abundant intraneuronal filamentous tau inclusions and extracellular filamentous deposits of amyloid-beta (1). Both types of inclusion form independently, but amyloid-beta deposits in the neocortex appear to be related to the formation of tau inclusions. Abundant tau inclusions form in the entorhinal cortex and the hippocampal formation before they appear in the neocortex, where they give rise to the symptoms of AD (2). This is compatible with the prion-like spreading of tau inclusions, where soluble monomers are recruited and assemble into filaments with the same structures as those of the seeds (3, 4). The prion concept is supported by studies using electron cryo-microscopy (cryo-EM) to determine the atomic structures of tau filaments from the brains of individuals with tauopathies (1, 5).

To study the mechanisms underlying the prion-like spreading of filaments with the Alzheimer tau fold, one has to develop systems of templated seeding that reproduce the atomic structures of the filament seeds from AD brains. This has recently been achieved *in vitro* using tau seeds from the brains of individuals with AD and recombinant tau with 12 phosphomimetic serine/threonine to aspartic acid substitutions (PAD12 tau) (6). In these studies, AD-seeded PAD12 tau filaments could be used as seeds in a second round of *in vitro* seeded assembly, while AD-seeded unmodified tau filaments were seeding-incompetent in a separate study (7). In adult wild-type mice, the intra-striatal injection of tau seeds extracted from the brains of individuals with AD (8) and corticobasal degeneration (9) has led to the seeded assembly of amyloid filaments made of 4-repeat (4R) mouse tau with the same structures as those of the seeds (10).

We now extend this work to human neurons in the form of cortical brain organoids. Previously, brain organoids from iPSCs of patients with familial AD were used to study the formation of amyloid-beta deposits and tau inclusions (11, 12). More recently, these studies were extended to vascularised neuroimmune organoids from human controls, which were exposed to AD brain extracts (13). However, none of those studies reported the presence of amyloid-beta and/or tau filaments. Here we show that when exposed to tau seeds extracted from the brains of individuals with AD, cortical organoids generated from iPSCs of human healthy subjects formed neurofibrillary tangles containing tau filaments with structures identical to those of paired helical filaments (PHFs).

## RESULTS

### Templated seeding by tau seeds from AD brains in human cortical brain organoids

Induced pluripotent stem cells (iPSCs) from healthy human subjects and isogenic iPSCs knockout for human *MAPT* were differentiated into cortical organoids containing neurons, astrocytes and oligodendrocytes as described (14). Tau seeds (500 ng) from two previously described cases of AD (15, 16) were added to the culture medium. At different times [3, 28, 92 and 129 days post-seeding (dps)] the organoids were fixed with paraformaldehyde or snap-frozen on dry ice. Immunohistochemistry of fixed organoids was performed with the conformational anti-tau antibody MC-1 (17). At 3 dps, MC-1 staining was restricted to the organoid surface, probably reflecting the presence of the tau seeds (Figure 1a). This staining had disappeared by day 28, when a small amount of MC-1 positivity was observed in the organoids. This staining had increased by 60 dps and was stronger and more diffuse at 92 and 129 dps (Figure 1a,b). These findings were quantified by estimating the percentage of MC-1 tau-positive area (Figure 1c). There was extensive co-localisation of MC-1 staining and staining with the amyloid dye pFTAA (18) (Figure S1). We also added different amounts of seed (4-2,000 ng) to the organoids and estimated the percentage of MC-1 tau-positive area at 129 dps (Figure 1d,e). Seeding activity plateaued at around 500-1,000 ng seed.

**Figure 1.**
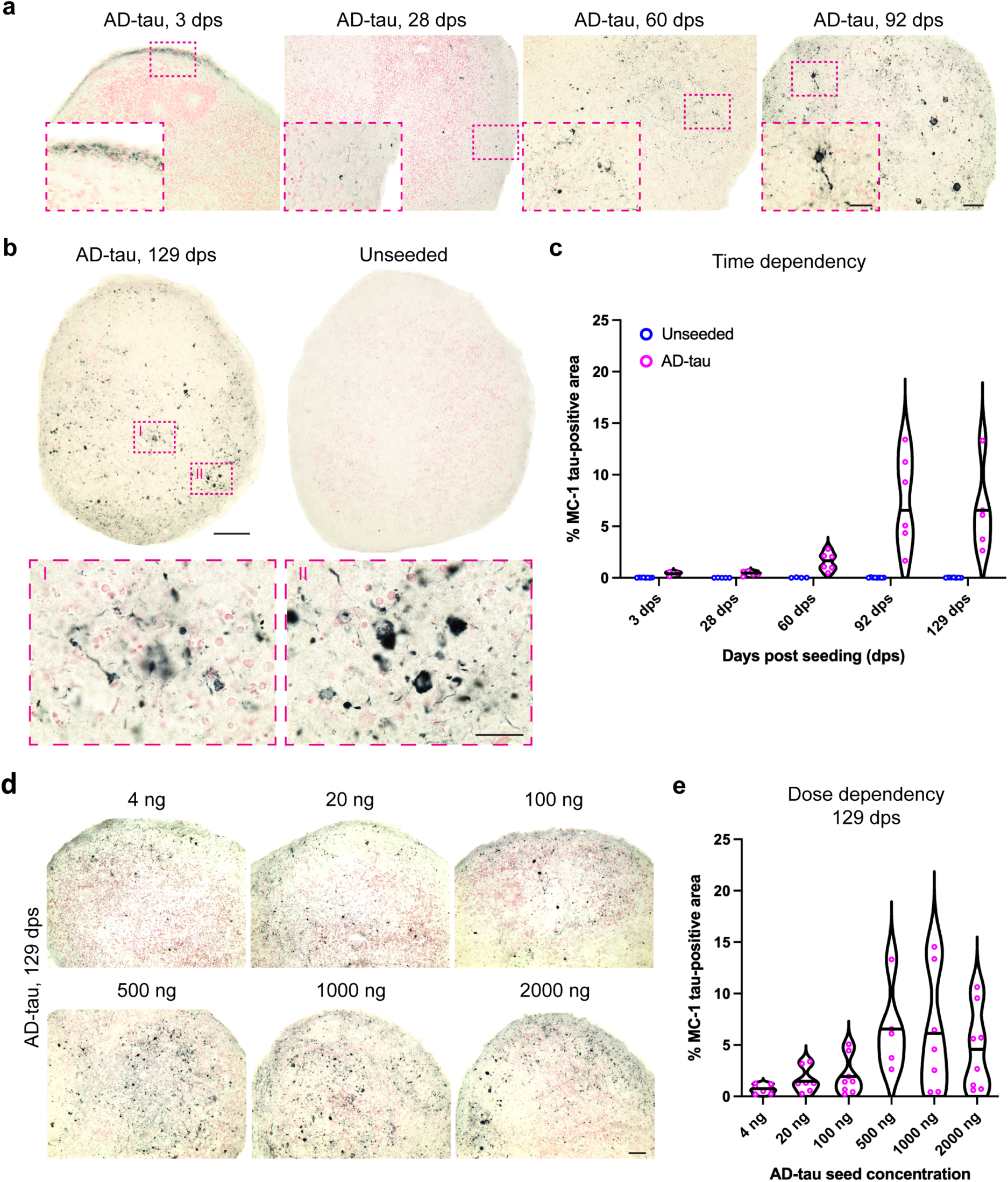
: Tau assembly in AD-tau seeded human cortical organoids is time- and concentration-dependent (a), Tau immunostaining of human cortical organoids with anti-tau antibody MC-1 at 3, 28, 60 and 92 days post-seeding (dps) using 500 ng AD-tau. Nuclear Fast red was used as counterstaining. Scale bars: 100 μm and 20 μm (insets). (b), Tau immunostaining of human cortical organoids with MC-1 at 129 dps using 500 ng AD-tau. Nuclear fast red was the counterstain. Insets (I and II) show higher magnifications of MC1-positive tau inclusions in seeded human cortical organoids. An unseeded organoid is also shown. Scale bars: 100 μm and 20 μm (insets). (c), Time dependency of tau assembly following seeding with 500 ng AD-tau (expressed as % MC-1 tau-positive area). Each dot represents a quantified human cortical organoid. Violin plots show the distribution of values, with horizontal black bars indicating mean values (n=3-8 organoids per time point). (d), Tau immunostaining of human cortical organoids with MC-1 at 129 dps following the addition of 4, 20, 100, 500, 1,000 and 2,000 ng AD-tau seeds. Tau levels were determined by dot blot analysis (Figure S3). Scale bar, 50 μm. (e), Dose dependency of tau assembly at 129 dps following seeding of human cortical organoids with 4-2,000 ng AD-tau (expressed as the percentage of MC-1 tau-positive area). Each dot represents a quantified human cortical organoid. Violin plots show the distribution of values, with horizontal black bars indicating mean values (n=5-8 organoids per time point). Organoids were generated from iPSC clone CRL1 (panels a-e).

### Templated seeding in human brain cortical organoids is dependent on the expression of tau

To determine the requirement of endogenous tau for templated seeding, we used the previously described exon 4-targeted isogenic *MAPT*^+/-^and *MAPT*^-/-^iPSC lines (19). They carry the same 25-bp deletion that leads to a frameshift and a stop codon within exon 4 of *MAPT*. A clone of the same iPSC line that failed to have its *MAPT* genetically edited served as an isogenic *MAPT*^+/+^ control. Figure 2a,b shows seeded assembly of tau in *MAPT*^+/+^, but not *MAPT*^+/-^or *MAPT*^-/-^human cortical organoids, as evidenced by immunostaining with MC-1. The same was shown by western blotting of sarkosyl-insoluble tau from extracted organoids using anti-tau antibody BR133 (Figure 2c). Templated seeding was dependent on the expression of human tau. Figure S2 illustrates that by western blotting with anti-tau antibody KJ9A that recognises all tau isoforms and antibody RD3 that is specific for 3-repeat (3R) tau, homogenates from *MAPT*^+/-^cells had lower levels of tau than *MAPT*^+/+^ cells, whereas no tau-positive bands were detected in *MAPT*^-/-^cells.

**Figure 2.**
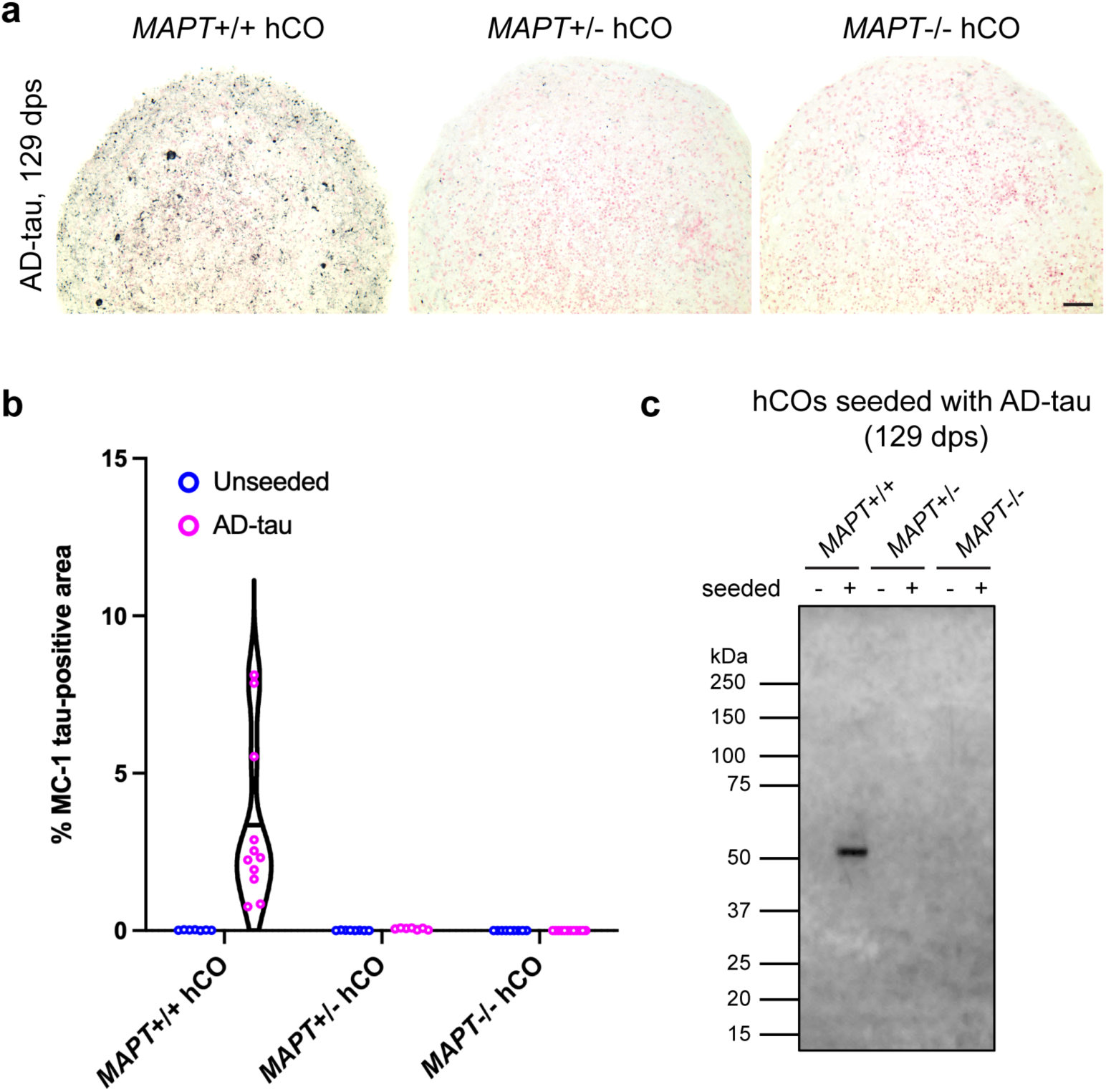
: Tau assembly in AD-tau seeded human cortical organoids is dependent on tau expression(a), Immunostaining of isogenic *MAPT*^+/+^, *MAPT*^+/-^and *MAPT*^-/-^human cortical organoids with MC-1 at 129 dps using 500 ng AD-tau. Nuclear fast red was used as the counterstain. (b), Tau assembly in isogenic *MAPT*^+/+^, *MAPT*^+/-^and *MAPT*^-/-^human cortical organoids at 129 dps with 500 ng AD-tau seeds (expressed as the percentage of MC-1 tau-positive area). Each dot represents one quantified human cortical organoid. Violin plots show the distribution of values, with horizontal black bars indicating mean values (n=7-12 organoids per time point). Unseeded organoids are also shown. (c), Sarkosyl-insoluble fractions from isogenic *MAPT*^+/+^, *MAPT*^+/-^and *MAPT*^-/-^human cortical organoids at 129 dps with 500 ng AD-tau seeds were immunoblotted with anti-tau antibody BR133.

### Tau seeds

Sarkosyl-insoluble tau seeds were isolated from the frontal cortex of two previously described cases of neuropathologically confirmed AD (15, 16). The cryo-EM structures of tau filaments from the second AD case are shown (Figure S3a,b), while those of the first case have been described previously (15); conversely, the characteristics of tau seeds from the second AD case have been described before (16), whereas those of the first case are shown here (Figure S3c,d). As expected (20), sarkosyl-insoluble tau from the first AD case consisted of three strong bands of 60, 64 and 68 kDa, and a weak band of 72 kDa (Figure S3c). By contrast, at 129 dps, seeded organoids had a single tau band of 55 kDa that was negative with RD4, an antibody specific for 4R tau (Figure S4).

### Seeded tau filaments in human brain cortical organoids and their cryo-EM structures

Transmission electron microscopy and immunoelectron microscopy were carried out on resin-embedded cortical organoids at 129 dps seeded with 500 ng AD-tau. The organoids contained abundant filaments that were decorated by anti-tau antibody MC-1 (Figure 3). Some filaments were in the form of parallel bundles (Figure 3b,f). We next determined the cryo-EM structures of templated tau assemblies at 129 dps (Figure 4). Following sarkosyl extraction, cryo-EM analysis showed that most tau filaments were PHFs identical to those from the brains of individuals with AD. More than 70% of filaments were made of two identical protofilaments with the Alzheimer tau fold, while 24% of filaments consisted of single protofilaments (Figure S5).

**Figure 3.**
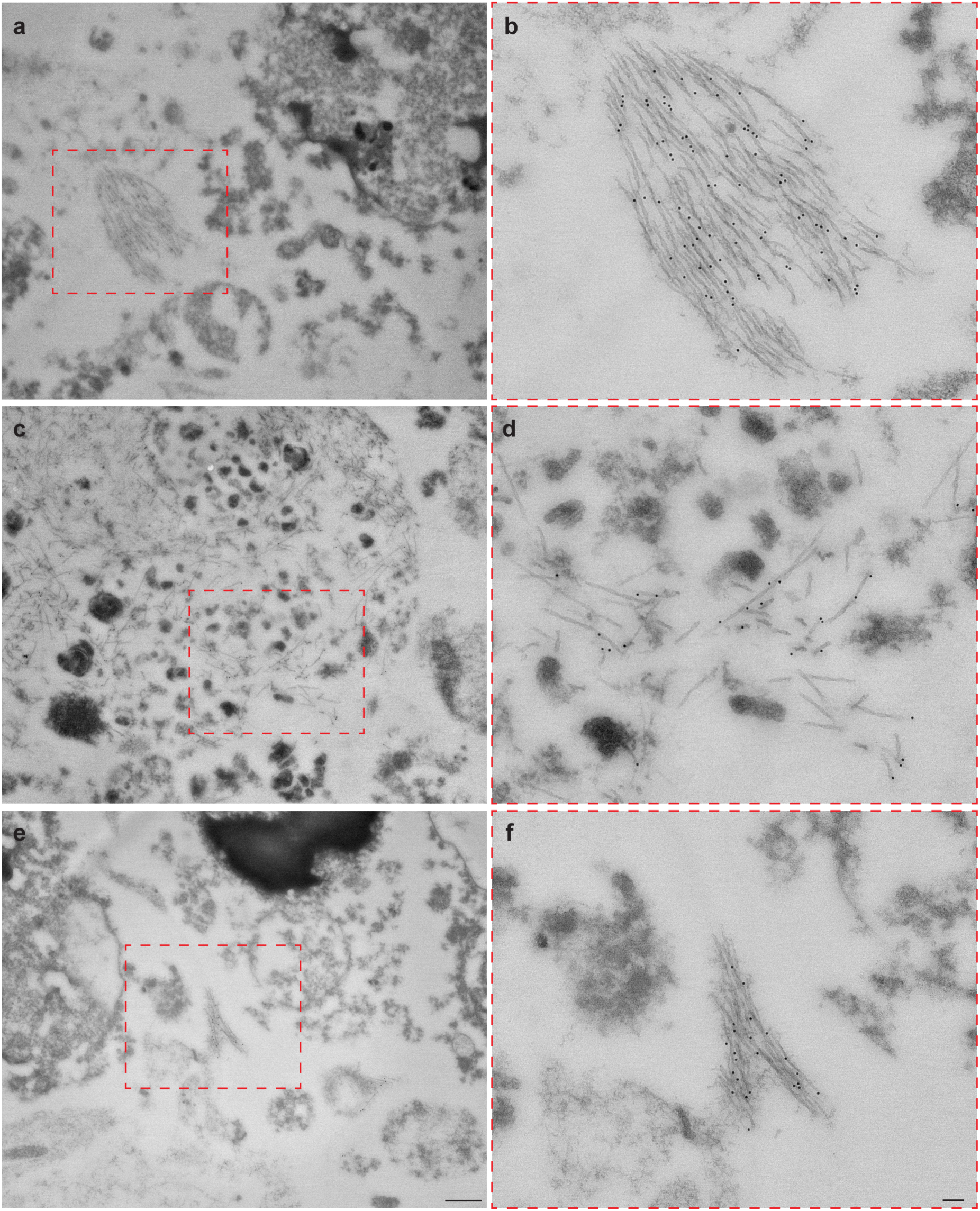
: Immunoelectron microscopy of tau filaments in AD-tau-seeded human cortical organoids (a,c,e) Representative transmission electron microscopy images of ultrathin sections from human cortical organoids at 129 dps with 500 ng AD-tau. Anti-tau antibody MC-1 was used. Scale bar in e (for a,c,e), 500 nm. Organoids were generated from tau iPSC clone CRL1. (b,d,f) Higher magnification images (red insets in a, c and e) show tau filaments decorated by gold particles that are conjugated with anti-tau antibody MC-1. Scale bar in f (for b,d,f), 100 nm.

**Figure 4.**
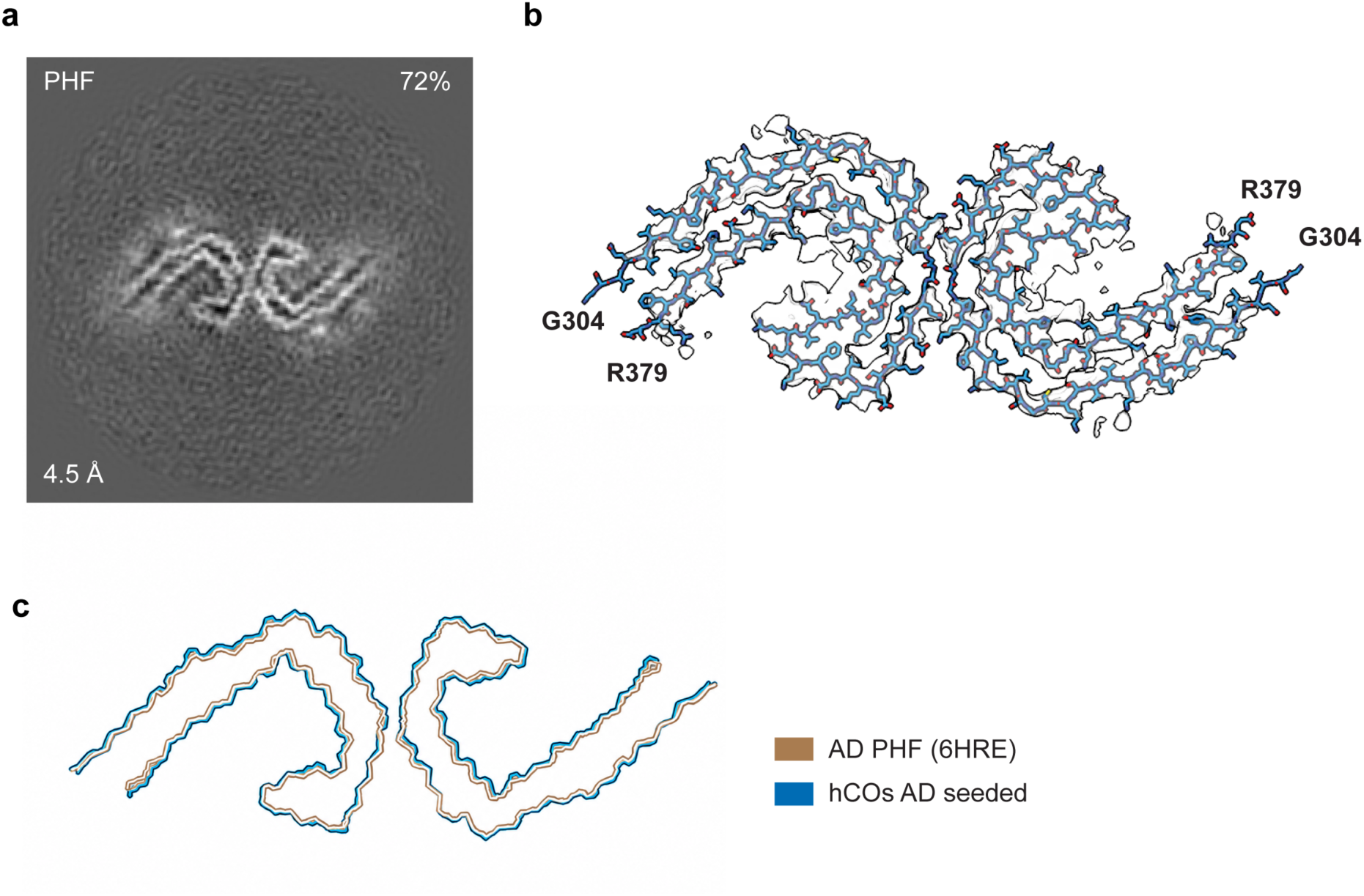
: Seeded filaments are identical to paired helical tau filaments extracted from AD brains (a), Cryo-EM cross-section of tau filament from seeded human cortical organoids at 129 dps. (b), Cryo-EM density map (in grey) and atomic model (in blue) of tau filament from seeded human cortical organoids at 129 dps. (c), Overlay of tau filament structures from seeded human cortical organoids at 129 dps (in blue) and the Alzheimer tau fold (in light brown) (PDB: 6HRE). Organoids were generated from iPSC clones CRL1, CRL2 and CRL3.

## DISCUSSION

The development of iPSC-derived brain organoids (21) has opened the way to studying human-specific aspects of neurodegeneration. Here we use these self-organising three-dimensional cells to investigate the structures of tau aggregates seeded with tau filaments extracted from the brains of individuals with AD. Much evidence indicates that tau filaments form in a limited number of brain cells, from where they spread in a prion-like manner. The prion concept relies on the formation of seeded aggregates having the same atomic structures as the seeds (4, 22). It is supported by our cryo-EM experiments which have shown that cases of primary age-related tauopathy at Braak stage II have the same tau filament structures in the entorhinal cortex as do cases of AD at Braak stage VI in the neocortex (8, 23). Tau filaments are believed to spread from the entorhinal cortex to the hippocampal formation and from there to the neocortex (2). There is no evidence to suggest that the structures of amyloid filaments change during the spreading of pathology.

Here we describe an experimental system in human brain cortical organoids for studying mechanisms of seeded aggregation. When seeds extracted from the frontal cortex of AD patients were added to the organoids, MC-1 immunoreactivity and pFTAA signal co-localised, indicating the formation of tau amyloid structures. Organoids expressing tau showed robust seeding, unlike those that were knockout for tau. Notably, isogenic heterozygous tau knockouts failed to show detectable seeded tau aggregation. This observation is consistent with seeded aggregation being sensitive to the levels of monomeric tau (1) Our results suggest that a lowering of physiological tau to around 50% is sufficient to prevent seeded aggregation and is thus informative for tau-lowering strategies, such as antisense oligonucleotides, as therapeutic avenues (24). Furthermore, the absence of detectable tau pathology in both the heterozygous and homozygous knockout organoids unambiguously establishes that expressed tau is necessary for aggregation, ruling out remaining seeds as the source of recovered tau aggregates.

Consistent with cortical organoids being at an early developmental stage, predominantly foetal human tau (0N3R) was expressed. The tau filaments extracted from AD brains and used to seed the cortical organoids contained both 3R and 4R tau (20). However, by western blotting the seeded tau filaments extracted from the organoids were negative with RD4, an antibody specific for 4R tau, excluding that the extracts obtained from the organoids contained the AD seeds. Even though AD-tau consists of 3R and 4R tau isoforms (20), the sequence of the core of the Alzheimer tau fold is shared by 3R and 4R tau (8). It follows that 3R and 4R tau alone can give rise to PHFs (6, 10).

Here we found that PHFs consisting of two identical 3R tau filaments with the Alzheimer fold form in seeded human cortical organoids. Over 70% of filaments were PHFs, with 24% being singlets. Previously, we showed that seeded aggregation with AD-tau in undifferentiated SH-SY5Y cells (25) and HEK293T cells (26) overexpressing tau failed to form PHFs, indicating the need for a nerve cell environment for forming PHFs. Interestingly, the Pick tau fold is made of 3R tau (27–29), here the filaments are made of 3R tau but have the structures of PHFs, suggesting that the structures of the seeds rather than the available tau isoforms determine the structures of the templated filaments.

It remains to be seen if seeds extracted from the brains of individuals with Pick’s disease can seed aggregation of the Pick tau fold in human brain organoids. The generation of human iPSCs making 4R tau by genome editing (30) has opened the way to test the seeded aggregation of 4R tauopathies in human brain organoids.

Based on studies in mice, it has been suggested that the transplantation of human nerve cells is necessary for studying the mechanisms of toxicity downstream of tau filament formation (31). The model described here has three major advantages over other experimental systems for human tauopathies. It forms PHFs like those in the AD brain, the cells are human neurons and they express endogenous levels of tau. Mechanisms involved in filament formation in human brain can be investigated here and the pathways involved can be manipulated to better understand the pathological process and identify new targets for therapy. Organoids also make it possible to study the effects of other cell types, such as astrocytesand microglia, on the mechanisms of toxicity (32). While they contain astrocytes and some oligodendrocytes, microglial cells are not present in our organoids, but they could be added to determine their contributions to tau aggregation and spreading as we have previously investigated in mouse primary neuronal and microglial co-cultures (33). Taken together, we describe a novel human system for studying the mechanisms of PHF formation and their downstream effects.

## MATERIALS and METHODS

### Maintenance of human iPSC lines

Human iPSC lines were derived from dermal fibroblasts of healthy donors with wildtype *MAPT*. Three clones [SIGi001-A (CRL1), CRL-2429 (CRL2) (34), and NN0003920 (CRL3); Table 1] were differentiated into organoids and gave similar results when seeded with AD-tau. In addition, isogenic iPSC lines from clone SFC856-03-04 were used that were either *MAPT*^+/+^, *MAPT*^+/-^or *MAPT*^-/-^(Table 1) (19). The iPSC lines were maintained on vitronectin (0.5 μg/ml, Gibco)-coated 6-well plates at 37° C with 5% CO_2_, with daily changes of mTeSR1 medium (Stem Cell Technology). The cells were passaged when they reached 80% confluency using ReLeSR (Stem Cell) and were plated in iPSC medium containing Rock inhibitor (10 mM, Y-27632 dihydrochloride, Tocris) for the first 24 h. For storage, following lifting with ReLeSR, confluent cells were frozen and stored in liquid nitrogen in 45% knockout serum replacement (Thermo Fisher/Gibco), 45% cell culture medium with Rock inhibitor and 10% DMSO.

**Table 1.** : Human iPSC lines

| **iPSC clones**<br>**(as referred in**<br>**the text)** | **iPSC clone ID** | **MAPT**<br>**Geno-**<br>**type** | **Sex** | **Age**<br>**(years)** | **Source/ Ref** |
| --- | --- | --- | --- | --- | --- |
| CRL1 | SIGi001-A | +/+ | F | 20-24 | Sigma-Aldrich |
| CRL2 | CRL-2429 | +/+ | M | newborn | ATCC, [34] |
| CRL3 | NN0003920 | +/+ | M | 64 | NINDS repository |
| MAPT+/+ line | SFC856-03-04 – B7 | +/+ | F | 78 | [19] |
| MAPT+/- line | SFC856-03-04 – A12 | +/- | F | 78 | [19] |
| MAPT-/- line | SFC856-03-04 – E1 | -/- | F | 78 | [19] |

### Differentiation of iPSCs into human cortical organoids

Cortical organoids were generated as described (14), with minor modifications. iPSCs were detached from the plates and dissociated into single-cell suspensions using ReLeSR and Accutase (Sigma Fine Chemicals). For embryoid body formation, 10,000 cells were plated per well of ultra-low attachment U-bottom 96-well plates (Corning) in iPSC medium plus Rock inhibitor. After 24 h, the medium was changed to neural induction medium (DMEM/F12, 20% knockout serum replacement, 1% MEM non-essential amino acids, 1% GlutaMAX, 1% penicillin/streptomycin, 100 μM 2-mercaptoethanol) supplemented with dual SMAD inhibitors dorsomorphin (5 μM, R&D Systems) and SB-421542 (10 μM Selleckchem) and changed daily. From days 6-24, human cortical organoids were fed with neural differentiation medium (Neurobasal A, B27 supplement without vitamin A, GlutaMax) supplemented with 20 ng/ml epidermal growth factor (EGF, Thermo Fisher) and 20 ng/ml fibroblast growth factor-2 (FGF-2, Prepro Tech). The medium was changed daily until day 16, from when it was changed every two days until day 25. From days 25-43, the neural differentiation medium was supplemented with 20 ng/ml brain-derived neurotrophic factor (BDNF, PreproTech) and 20 ng/ml neurotrophin-3 (NT3, PreproTech) and changed every two days. The human cortical organoids were then maintained in neural differentiation medium.

### Extraction of tau filament seeds from AD brains

Sarkosyl-insoluble material was extracted from the frontal cortex of two previously described cases of sporadic AD (15, 16), as described (35, 36). In (15), AD case 1 corresponds to case 2. The pelleted sarkosyl-insoluble material was resuspended in 10 mM Tris-HCl, pH 7.4, 800 mM NaCl, 5 mM EDTA and 1 mM EGTA, with a final concentration of 10% (w/v) sucrose at 750 μl/g tissue, followed by centrifugation at 20,100 g for 30 min at 4° C. The pellets were resuspended in 20 mM Tris-HCl, pH 7.4, containing 100mM NaCl at 250 μl/g tissue and centrifuged at 100,000 g for 30 min at 4° C. The final pellets were resuspended in buffer at 50 μl/g frontal cortex.

### Dot blot analysis of AD-tau seeds

Tau concentrations in AD extracts were measured using a quantitative filter retardation assay (26). Briefly, sarkosyl-insoluble fractions from AD case 1 (15) were applied in triplicate to 0.2 μm nitrocellulose membranes (Bio-Rad) using a Bio-Dot microfiltration apparatus (Bio-Rad). In parallel, tau filaments of known concentration that were formed by incubating 2N4R recombinant tau with heparin were also applied. The membranes were blocked with 3% bovine serum albumin (Thermo Fisher) for 1h at room temperature, before an overnight incubation at 4° C with anti-tau antibody BR134 (1:3,000). The membranes were then washed three times in phosphate-buffered saline + 0.3% Triton-X100 (PBST) and incubated for 1h at room temperature with Alexa-conjugated secondary antibodies (1:2,000). They were imaged using a ChemiDoc gel imager system (Bio-Rad) and the dots were quantified using Fiji software (36). The quantification of sarkosyl-insoluble fractions from AD case 2 has been described (16).

### Seeding in human cortical organoids with AD brain-derived tau filaments

AD tau seeds were added once to the culture medium of human cortical organoids aged 2-3 months (typically 500 ng seed). At 3 dps the medium was changed to neural differentiation medium. The organoids were cultured in this medium until harvesting, with full medium changes every 3-4 days.

### Histochemistry

Organoids were collected at different time points, washed once in PBS and fixed with 4% paraformaldehyde for 1h at room temperature, followed by three more washes. They were then transferred to a 30% sucrose solution overnight at 4° C. Organoids were next embedded in 7.5% gelatine/10% sucrose/0.05% NaN_3_ in PBS and snap frozen in cold isopentane on dry ice, before being kept at -80° C until sectioning. Sections of 15 μm were cut on a cryostat and stored at -20° C until further use. For immunohistochemistry, sections were rehydrated in PBS for 5 min and endogenous peroxidase was quenched by a 30 min incubation in 3% hydrogen peroxide/20% methanol in PBS. Following a brief wash in PBST, the sections were incubated in blocking buffer (2.5% bovine serum albumin, 2.5% horse serum in PBST) for 1h at room temperature. They were then incubated with the conformation-dependent anti-tau antibody MC-1 (1:500) (17) overnight at 4° C. Following washing in PBS, the sections were incubated with ImmPRESS-HRP polymer anti-mouse detection antibodies (Vector Laboratories) for 2h at room temperature. Following washing with PBS, ImmPACT vector SG peroxidase substrate (Vector Laboratories) was added to visualise the signal. The sections were counterstained with nuclear fast red (Vector Laboratories) for 2 min at room temperature. Following dehydration, the sections were coverslipped with Entellan mounting medium (Merck). Images were acquired using a QImaging Retiga 2,000 CCD camera mounted on an Olympus BX50 microscope or a slide scanner (Zeiss Axio Scan.Z1, 40X magnification).

For fluorescence, following an overnight incubation at 4° C with MC-1 (1:500) and a wash in PBS, the sections were incubated with Alexa Fluor 647-conjugated secondary antibody (1:400, Invitrogen) and with 3 μM of pFTAA (18) in PBS, including 1 μg/ml Hoechst dye 33342 (Sigma Fine Chemicals) for 2 h at room temperature. Following washing in PBS, the sections were treated with TrueBlack lipofuscin autofluorescence quencher (Biotum) for 2 min at room temperature. After washing in PBS, they were coverslipped with Fluoromount-G mounting medium (Southern Biotech) and stored at 4° C. Images were captured on a confocal microscope (Leica Stellaris). For image modification and quantification of the stained areas, FIJI (-37) or QuPath (38) software was used. Brightness and contrast were adjusted for optimal signal visualisation.

### Immunogold electron microscopy of organoid sections

Following fixation in 4% paraformaldehyde at room temperature for 1h, human cortical organoids were washed four times in 0.05M sodium cacodylate buffer (pH 7.4) and once in water. The samples were then dehydrated in an ascending ethanol series, before being infiltrated overnight at 4° C with a 50:50 mixture of 100% ethanol and London Resin White (LRW). The following day, LRW was exchanged three times and one more time the day after. The organoids were then embedded in gelatine capsules, covered with Aclar coverslips and the resin was cured for 24 h at 55° C in an embedding oven. Sections were cut using a Leica Ultracut ultramicrotome at a thickness of 80 nm; they were placed on bare copper grids for transmission electron microscopy and nickel-formvar grids for immune-transmission electron microscopy. Sections were then allowed to air dry. For immunostaining, the grids were floated on blocking solution (0.1% bovine serum albumin/0.01% Tween-20 in PBS) for 30 min at room temperature, followed by an overnight incubation at 4° C with anti-tau antibody MC-1 (1:10 in blocking solution). Excess primary antibody was removed with PBST. The grids were then floated on goat anti-mouse secondary antibodies conjugated to 10 nm gold particles (1:20; Abcam) in blocking solution for 1h at room temperature. Excess secondary antibody was removed with PBST. The sections were post-stained using 2% aqueous uranyl acetate (3 min) and Reynold’s lead citrate (6 min). Images were captured on a Tecnai G2 transmission electron microscope (Thermo Fisher Scientific) run at 200 keV using a 20 µm objective aperture to improve contrast. Images were acquired using an ORCA HR high-resolution CCD camera (Advanced Microscopy Techniques).

### Extraction of seeded tau filaments from human cortical organoids

Sarkosyl-insoluble material was extracted from 10-30 pooled human cortical organoids, as described (36), with minor modifications. Briefly, organoids were added to 75 μl buffer A/organoid [10 mM Tris-HCl, pH 7.4, 800 mM NaCl, 10% sucrose, 1 mM EGTA, supplemented with cOmplete EDTA-free protease inhibitor cocktail, PhosSTOP phosphatase inhibitor cocktail (Roche) and 1% sarkosyl]. Following sonication with a Microson XI-2000 Ultrasonic cell disruptor (Miso-nix) for 15 s and a 30 min incubation at 37° C while shaking, total lysates were cleared by centrifugation (10,000 g for 10 min at 4° C). The supernatants were then spun at 150,000 g for 1h at 4° C. The pellets were resuspended in 25 μl buffer A/organoid, diluted 3-fold with buffer B (50 mM Tris-HCl, pH 7.5, 150 mM NaCl, 10% sucrose, 0.2% sarkosyl) and spun again at 150,000 g for 1h at 4° C. The final pellets were resuspended in 1.5 μl/organoid of 50 mM Tris-HCl, pH 7.4, 150 mM NaCl.

### Western blotting

One μl of sarkosyl-insoluble material from organoids (1.5 μl/organoid) was loaded per lane. For AD-tau seeds, 0.3 μl of sarkosyl-insoluble material (50 μl/g frontal cortex) was loaded. For total lysates, 6 μl were loaded per lane. NuPAGE Bis-Tris 4-12% gels (Thermo Fisher Scientific) were electroblotted onto 0.2 μm nitrocellulose membranes using the transblot turbo transfer system (Bio-Rad). The membranes were blocked in 3% bovine serum albumin in PBST for 1 h at room temperature, followed by an overnight incubation at 4° C with primary anti-tau antibodies BR133 [1:3,000; (39)], BR134 [1:3,000; (38)], K9JA (1:5,000; Agilent), RD3 (1:3,000; Millipore)and RD4 (1:3,000; Millipore), as well as a GAPDH antibody (1:3,000; Thermo Fisher Scientific). BR133, BR134 and K9JA label all six human brain tau isoforms. RD3 is specific for 3R tau and RD4 is specific for 4R tau. Following repeated washes with PBST, the membranes were incubated with Alexa-conjugated secondary antibodies (A-11008, A21244, A-21235, A-21121; Thermo Fisher Scientific) for 1h at room temperature. Signal was imaged using a ChemiDoc gel imager (Bio-Rad).

### Cryo-EM sample preparation and data collection

Cryo-EM grids (Quantifoil Au R1.2/1.3, 300 mesh) were glow-discharged for 1 min using an Edwards (S150b) sputter coater. For the tau seeds from the second AD case, 2 μl of the resuspended final pellet were diluted 1:6 and 3 μl of the diluted sample were applied to each glow-discharged grid. For the seeded tau aggregates from the human cortical organoids, 3 μl of the final pellets were applied to each glow-discharged grid. The grids were blotted with filter paper (blot force 20, wait time 0s, drain time 0s, blot time 4s) and plunge-frozen into liquid ethane using Vitrobot Mark IV (FEI) (100% humidity and 4° C). Cryo-EM images were acquired with a Titan Krios microscope using EPU software (Thermo Fisher Scientific) operated at 300 kV with a total dose of 40 electrons/A^2^. For AD case 2, movies were acquired using a Gatan K3 detector at a pixel size of 0.826 Å/pixel and an energy filter with a slit width of 20 eV. For the human cortical organoids, movies were acquired using a Falcon-4i detector at a pixel size of 0.824 Å/pixel. See Table S1 for further details.

### Cryo-EM data processing

Datasets were processed in RELION using standard amyloid helical procedures (40). Movie frames were gain-corrected, aligned and dose-weighted using RELION’s own motion correction program (41). Contrast transfer functions (CTFs) were estimated using CTFFIND 4.1 (42). Tau filaments from seeded human cortical organoids were picked manually, while filaments from AD case 2 were picked automatically (43). Particles were initially extracted in box sizes of 768 or 1,024 pixels and downscaled to 256 pixels. Reference-free 2D classification was used to distinguish between different filament types, discard suboptimal particles and measure cross-over distances for initial model calculation using relion_helix_inimodel2d (44). Particles for subsequent three-dimensional auto-refinements were extracted in a box size of 384 pixels, auto-refinements were performed with optimisation of the helical twist and rise parameters. Bayesian polishing (45) and CTF refinement (41) were used to increase the resolution of the final map, which was sharpened using post-processing procedures in RELION. Final resolution estimates were calculated based on Fourier shell correlations (FSC) between two independently refined half-maps using the 0.143 criterion (46).

### Model building and refinement

The atomic model (PDB: 6HRE) (8) was rigid-body fitted into the cryo-EM density of the organoid-seeded PHFs using ISOLDE (47, 48), without further atomic refinement due to the limited resolution of the cryo-EM density (4.5 Å). Figures were prepared with ChimeraX (49).

## List of Supplementary Materials

Fig. S1: MC-1 and pFTAA double-labelling of seeded human cortical organoids at 129 days post-seeding.

Fig. S2: Tau expression in lysates of human cortical organoids.

Fig. S3: Characterisation of sarkosyl-insoluble tau seeds from AD cases 1 and 2.

Fig. S4: Western blotting of seeded tau filaments.

Fig. S5: Cryo-EM structure determination of tau filaments from seeded human cortical organoids.

Table S1: Cryo-EM data acquisition and maps refinement details.

## Acknowledgements

We thank the participants’ families for donating brain tissues, T. Darling, I. Clayson and J. Grimmett for help with high-performance computing and the EM facility of the Medical Research Council (MRC) Laboratory of Molecular Biology for help with cryo-EM data acquisition. Immuno-EM studies were performed at the EM-facility of the Microscopy Bioscience Platform, Cambridge. We thank L. Bedwell, L. Butter and Z. Zheng for help with sectioning of organoids. M.G. and M.G.S. are Affiliate Members of the UK Dementia Research Institute.

## Author contributions

M.B. and M.G.S. designed the experiments and the overall study. M.B. generated hCOs and with J.B. and R.C. maintained hCOs. M.B., M.S. and T.K. extracted hCOs and patient-derived brain fractions. T.K. and W.A.M. provided tau seeds. M.B. performed histological experiments, and together with R.C. and J.B acquired the histology images.

M.S. and T.K. performed cryo-EM experiments. S.L. and S.H.W.S. contributed to the cryo-EM data analysis. F.G. and K.H.M. performed electron microscopy in organoid sections. S.A.C. provided *MAPT* iPSC lines. M.B., T.K., M.G. and M.G.S wrote the manuscript and all authors provided comments and approved the submission.

## Copyright statement

For the purpose of open access, the MRC Laboratory of Molecular Biology has applied a CC BY public copyright licence to any Autor Accepted Manuscript version arising.

