## Supplementary Material for "Templated seeding induces the formation of Alzheimer paired helical tau filaments in human brain organoids"

Bacioglu *et al.*

Corresponding author: Maria Grazia Spillantini,

**This PDF includes:**

Figs S1 to S5

Table S1

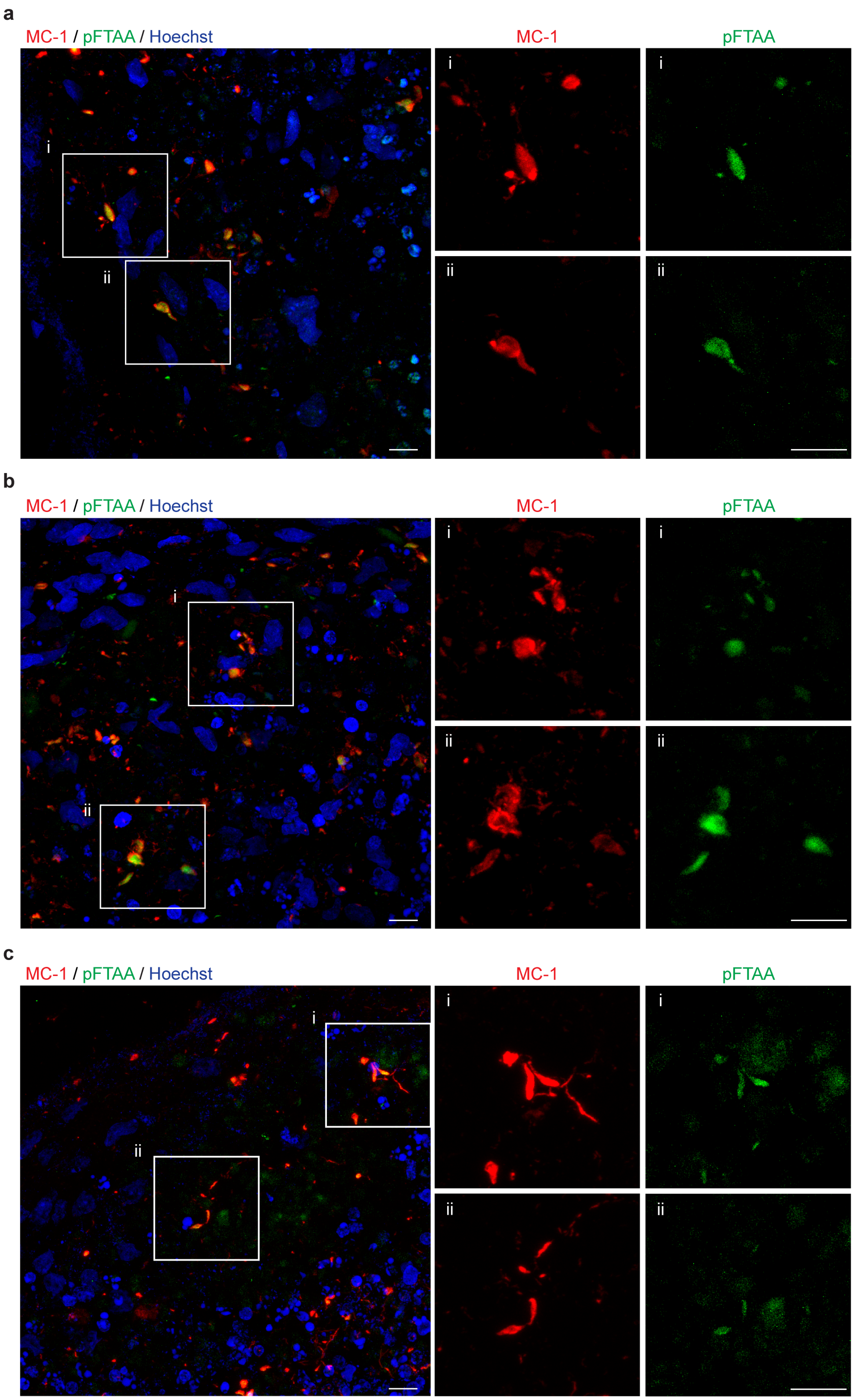
a
MC-1 / pFTAA / Hoechst
MC-1
pFTAA
i
ii
b
MC-1 / pFTAA / Hoechst
MC-1
pFTAA
i
ii
c
MC-1 / pFTAA / Hoechst
MC-1
pFTAA
i
ii

**Figure S1**

**MC-1 and pFTAA double-labelling of seeded human cortical organoids at 129 days post-seeding.**

Double-labelling fluorescence. MC-1 immunostaining is shown in red and pFTAA staining in green.

(i,i), The insets show MC-1 immunofluorescence (Red).

(ii,ii), The insets show pFTAA fluorescence (Green).

Organoids in (a,b) are from CRL 1 iPSC line.

Organoids in (c) are from *MAPT*$^{+/+}$ iPSC line. Scale bars, 10 μm.

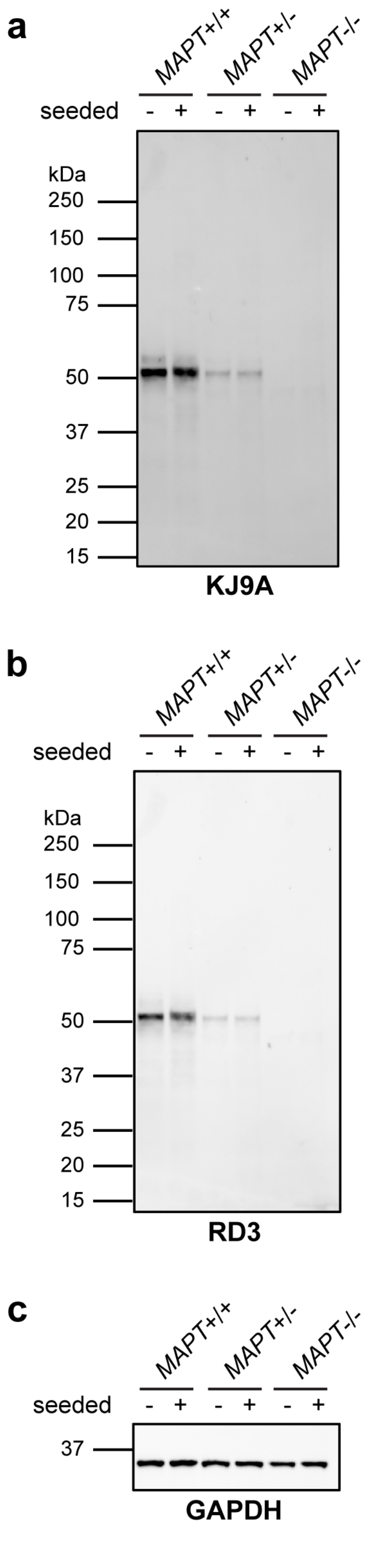
a
MAPT+/+
MAPT+/-
MAPT-/-
seeded
- + - + - +
kDa
250
150
100
75
50
37
25
20
15
KJ9A
b
MAPT+/+
MAPT+/-
MAPT-/-
seeded
- + - + - +
kDa
250
150
100
75
50
37
25
20
15
RD3
c
MAPT+/+
MAPT+/-
MAPT-/-
seeded
- + - + - +
37
GAPDH

**Figure S2**

**Tau expression in lysates of human cortical organoids.**

Western blot analysis of total lysates from isogenic *MAPT*$^{+/+}$, *MAPT*$^{+/-}$ and *MAPT*$^{-/-}$ human brain cortical organoids using anti-tau antibodies KJ9A (a) and RD3 (b), as well as a GAPDH antibody (c) as the loading control.

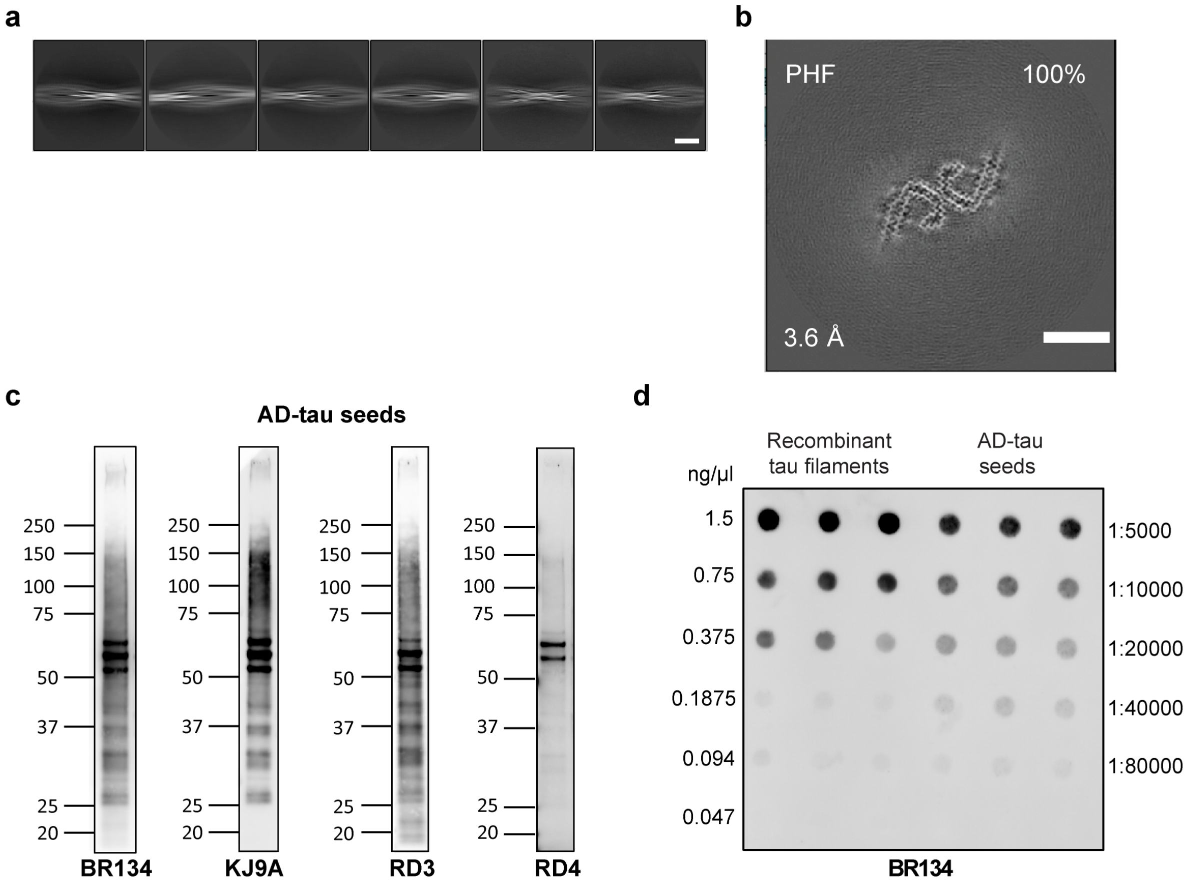

**Figure S3**

**Characterisation of sarkosyl-insoluble tau seeds from AD cases 1 and 2.**

(a), Representative 2D-classification images of sarkosyl-insoluble filaments extracted from the frontal cortex of AD case 1. Scale bar, 20 nm.

(b), Cryo-EM cross-section of tau filament from AD case 2 perpendicular to the helical axis, representing approximately one rung in projected thickness. The structural resolution (in Å) and percentage of filament type (%) are indicated at the bottom left and top right, respectively. Scale bar, 10 nm.

(c), Western blot analysis of tau seeds from AD case 1 using anti-tau antibodies BR134, KJ9A, RD3 and RD4.

(d), Dot blot showing estimation of tau levels in seeds from AD case 1.

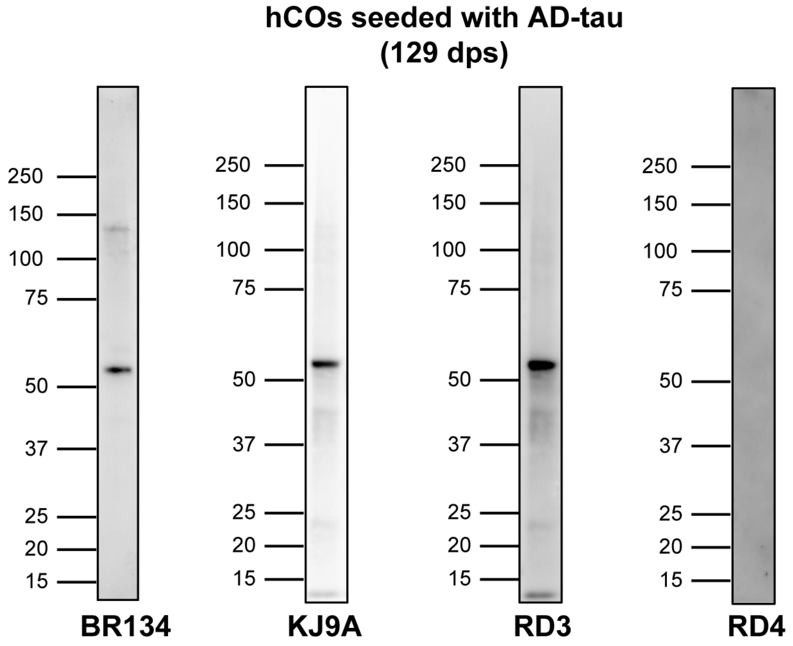


**Figure S4**

**Western blotting of seeded tau filaments.**

Sarkosyl-insoluble fractions from pooled (n=12) human cortical organoids at 129 dps with 500 ng AD-tau seeds were immunoblotted with anti-tau antibodies BR134, KJ9A, RD3 and RD4. Organoids were generated from iPSC clone CRL2.

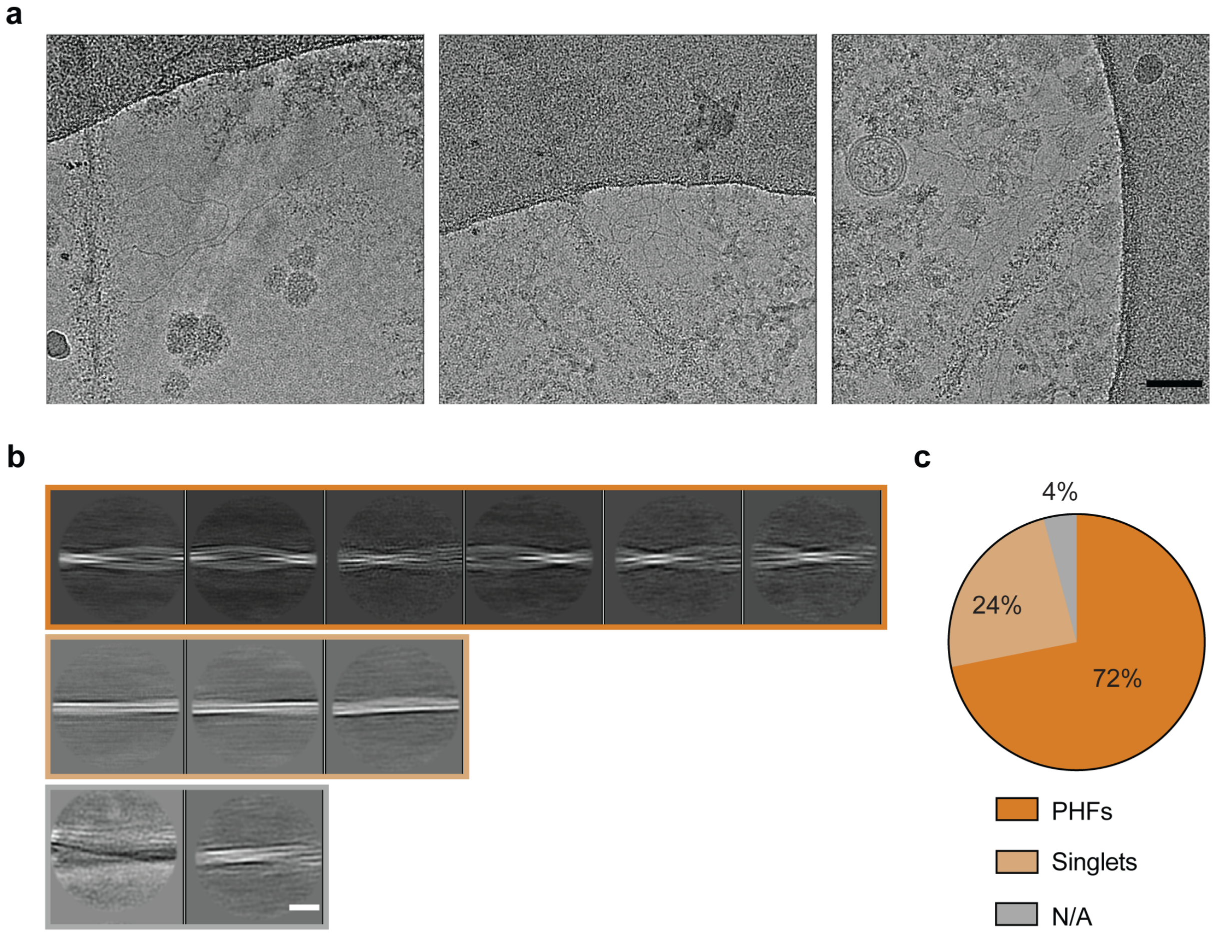


**Figure S5**

**Cryo-EM structure determination of tau filaments from seeded human cortical organoids.**

(a), Cryo-EM micrographs of tau filaments from seeded human cortical organoids. Scale bar, 20 nm.

(b), Representative 2D classification images of filaments from AD-seeded human cortical organoids. Scale bar, 20 nm.

(c), Pie chart with the particle distribution per filament type [two identical protofilaments with the Alzheimer tau fold in 72% of filaments (dark orange); a single protofilament with the Alzheimer tau fold in 24% of filaments (light orange); discarded filaments in 4% (grey).

**Table S1. Cryo-EM data acquisition and maps refinement details**

| | AD Case #2 | hCOS AD-seeded |
|---|---|---|
| **Data acquisition** | | |
| Pixel size (Å) | 0.826 | 0.824 |
| Voltage (kV) | 300 | 300 |
| Detector | K3 | Falcon 4i |
| Electron dose (e–/Å$^2$) | 40.0 | 40.0 |
| Energy filter slit (eV) | 20 | N/A |
| Defocus range (μm) | -1.0 to -2.0 | -1.0 to -2.0 |
| **Map refinement** | | |
| Symmetry imposed | C1 | C1 |
| Initial particle images (no.) | 3,386,690 (automatic) | 36,561 (Manual) |
| Final particle images (no.) | 379,384 | 26,279 |
| Symmetry imposed | C1 | C1 |
| Map resolution (Å) FSC threshold = 0.143 | 3.60 | 4.47 |
| Helical rise (Å) | 2.37 | 2.37 |
| Helical twist (°) | 179.45 | 179.45 |
